# Tau filaments with the Alzheimer fold in cases with *MAPT* mutations V337M and R406W

**DOI:** 10.1101/2024.04.29.591661

**Authors:** Chao Qi, Sofia Lövestam, Alexey G. Murzin, Sew Peak-Chew, Catarina Franco, Marika Bogdani, Caitlin Latimer, Jill R. Murrell, Patrick W. Cullinane, Zane Jaunmuktane, Thomas D. Bird, Bernardino Ghetti, Sjors H.W. Scheres, Michel Goedert

## Abstract

Frontotemporal dementia (FTD) and Alzheimer’s disease are the most common forms of early-onset dementia. Dominantly inherited mutations in *MAPT*, the microtubule-associated protein tau gene, cause FTD and parkinsonism linked to chromosome 17 (FTDP-17). Individuals with FTDP-17 develop abundant filamentous tau inclusions in brain cells. Here we used electron cryo-microscopy to determine the structures of tau filaments from the brains of individuals with *MAPT* mutations V337M and R406W. Both mutations gave rise to tau filaments with the Alzheimer fold, which consisted of paired helical filaments in all V337M and R406W cases and of straight filaments in two V337M cases. We also identified a new assembly of the Alzheimer fold into triple tau filaments in a V337M case. Filaments assembled from recombinant tau(297-391) with mutation V337M had the Alzheimer fold and showed an increased rate of assembly.

---

In the adult human brain, six tau isoforms are expressed from a single gene by alternative mRNA splicing (1). They differ by the presence or absence of one or two inserts in the N-terminal half and an insert in the C-terminal half. The latter encodes a repeat of 31 amino acids, giving rise to three isoforms with four repeats (4R). The other three isoforms have three repeats (3R). Together with adjoining sequences, these repeats constitute the microtubule-binding domains of tau (2). Some of the repeats also form the cores of assembled tau in neurodegenerative diseases, suggesting that physiological function and pathological assembly are mutually exclusive. Most *MAPT* mutations are in exons 9-12, which encode R1-R4, with some mutations being present in exon 13, which encodes the sequence from the end of R4 to the C-terminus of tau. Only two of the sixty-five known *MAPT* mutations are found near the N-terminus of tau (3). Mutations in *MAPT* lead to the formation of filamentous inclusions that are made of either 3R, 4R or 3R+4R tau (4). Mutations that cause the relative overproduction of wild-type 3R or 4R tau result in the deposition of 3R tau with the Pick fold (5) or 4R tau with the argyrophilic grain disease (AGD) fold (6). In cases of sporadic and familial tauopathies, filaments of TMEM106B also form in an age-related manner (7–9).

Structures of 3R+4R tau-containing filaments from cases with *MAPT* mutations have not been reported. In sporadic diseases, filaments made of 3R+4R tau have the Alzheimer (10) or the chronic traumatic encephalopathy (CTE) (11) fold. The Alzheimer tau fold is also found in familial British and Danish dementias, cases of prion protein amyloidoses and primary age-related tauopathy (PART) (6,12). The CTE tau fold is also typical of subacute sclerosing panencephalitis, amyotrophic lateral sclerosis/parkinsonism dementia complex and vacuolar tauopathy (13–15). Recombinant tau(297-391) forms filaments with either fold, depending on the *in vitro* assembly conditions (16).

Dominantly inherited mutations V337M (17–21) in exon 12 and R406W (22–28) in exon 13 of *MAPT* give rise to FTD with inclusions that are also made of all six tau isoforms (18,24). Mutation V337M, which is located inside the ordered cores of tau filaments (4), causes behavioural-variant FTD and cognitive impairment in the fifth or sixth decade (17,20,29); it has been reported that tau inclusions are abundant in cerebral cortex, but not in hippocampus (17). Mutation R406W, which is located outside the ordered cores of tau filaments, is associated with an Alzheimer’s disease (AD)-like amnestic phenotype that is characterised by initial memory impairment (30,31); abundant tau inclusions are present in both cerebral cortex and hippocampus (22). Here, we show that tau filaments from the brains of individuals with mutations V337M and R406W in *MAPT* adopt the Alzheimer fold.

## RESULTS

### Structures of tau filaments from three cases of Seattle family A with mutation V337M in *MAPT*

We used electron cryo-microscopy (cryo-EM) to determine the atomic structures of tau filaments from the frontal cortex of three previously described individuals of Seattle family A with mutation V337M in *MAPT* (Figures 1 and 2) (17,20). By immunohistochemistry, abundant tau inclusions were present in the frontal cortex (Extended Data Figure 1A) (17,18). Unlike previous reports (17), we detected hippocampal tau inclusions (Extended Data Figure 1B).

**Figure 1.**
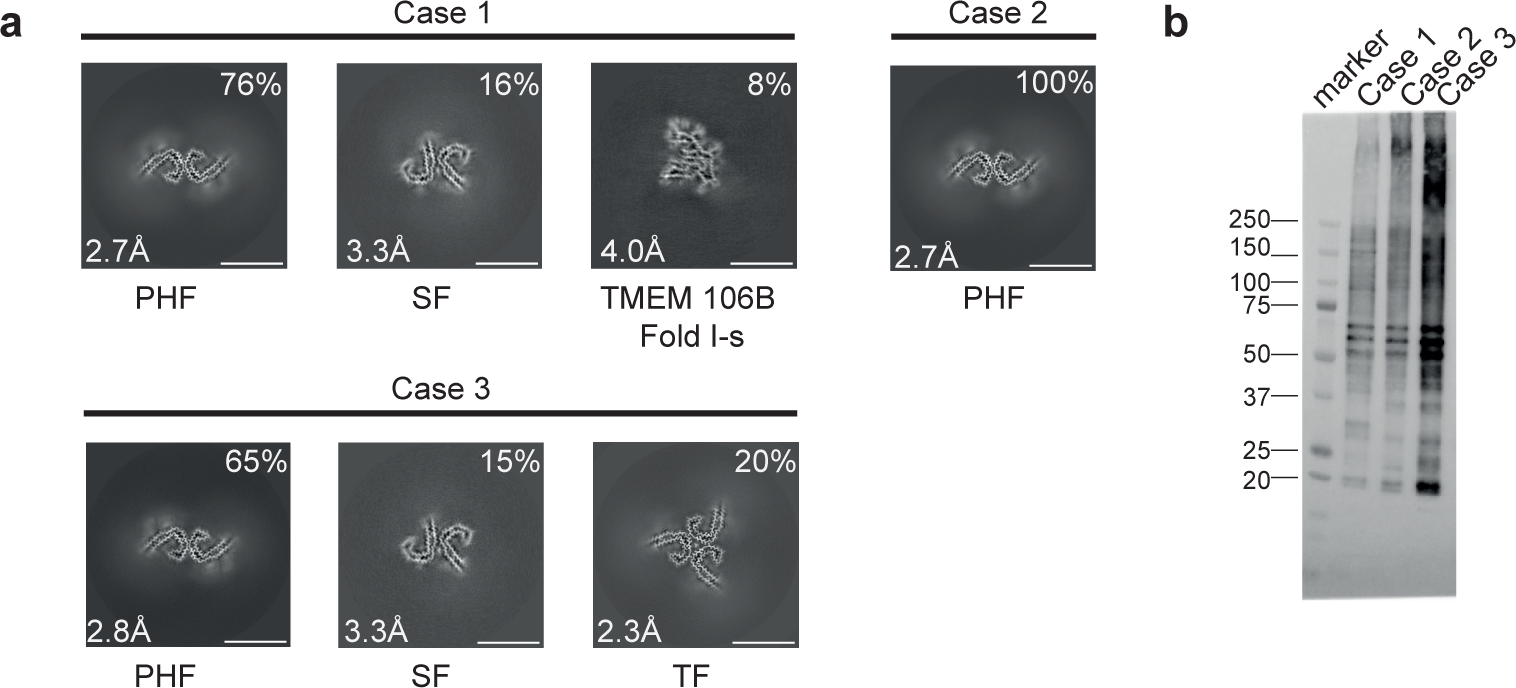
V337M mutation in *MAPT*: Cryo-EM cross-sections of tau filaments and immunolabelling. a, Cross-sections through the cryo-EM reconstructions, perpendicular to the helical axis and with a projected thickness of approximately one rung, are shown for the frontal cortex from cases 1-3. Resolutions (in Å) and percentages of filament types are indicated at the bottom left and top right, respectively. Scale bar, 10 nm. b, Immunoblotting of sarkosyl-insoluble tau from the frontal cortex of cases 1-3 with mutation V337M. Phosphorylation-independent anti-tau antibody BR134 was used.

**Figure 2.**
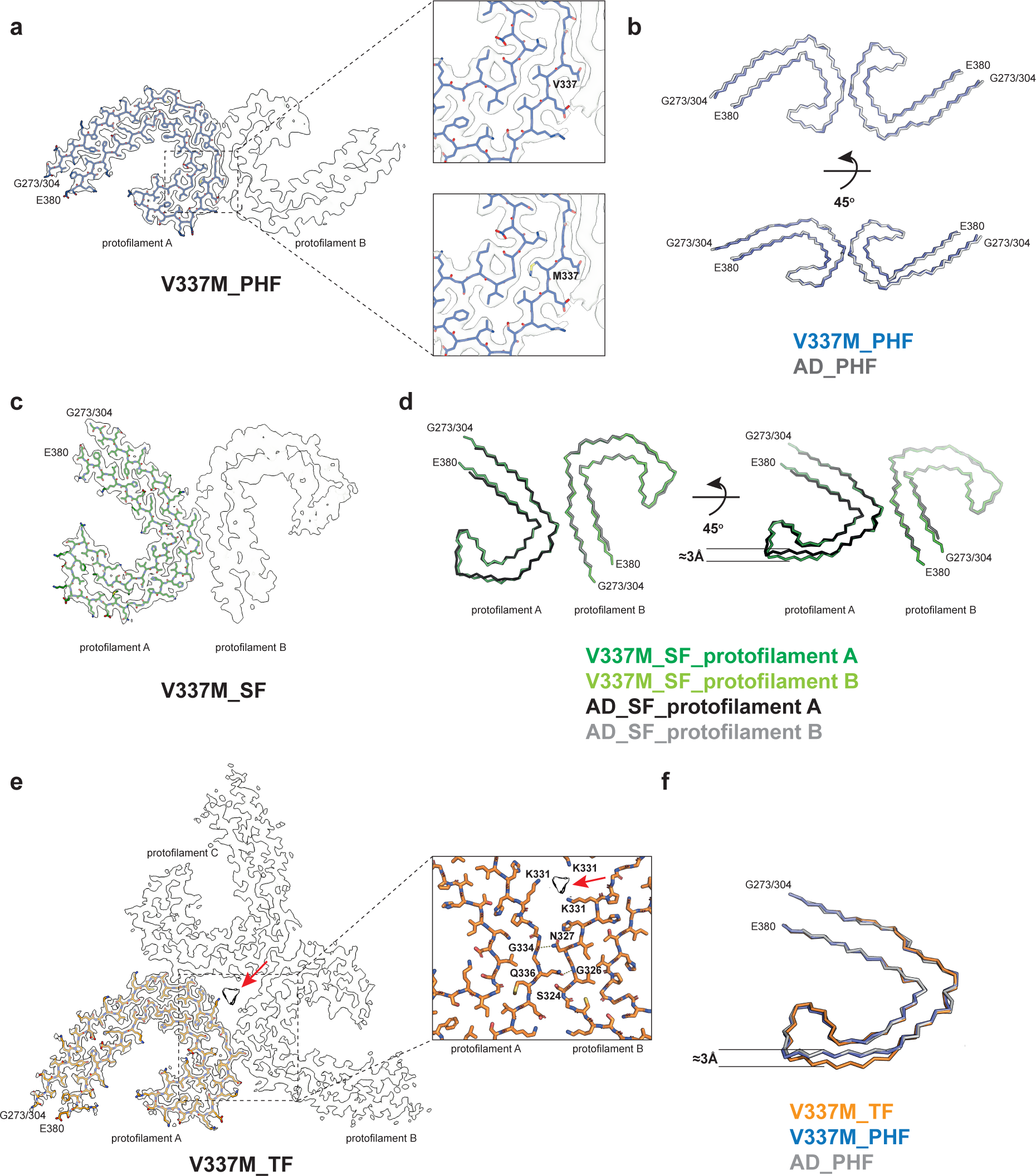
V337M mutation in *MAPT*: Cryo-EM structures of tau filaments. a, Cryo-EM density map and atomic model of paired helical filament (PHF). Two identical protofilaments extend from G273/304-E380. Inset: Zoomed-in view showing that both wild-type (V) and mutant (M) residues can fit into the density at position 337. b, Backbone representation of overlay of PHF extracted from the frontal cortex of case 1 with mutation V337M in *MAPT* (blue) and PHF extracted from the frontal cortex of an individual with sporadic AD (white, PDB:5O3L). The root mean square deviation (rmsd) between Cα atoms of the two structures is 0.78 Å. c, Cryo-EM density map and atomic model of straight filament (SF). Two asymmetrically packed protofilaments A and B extend from G273/304-380. d, Overlay of SF extracted from the frontal cortex of case 1 with mutation V337M (protofilament A is in dark green and protofilament B is in light green) and SF extracted from the frontal cortex of an individual with AD (PDB:5O3T) (protofilament A is in black and protofilament B is in grey). In protofilament A, strand β4 (residues 336-341) is shifted along the helical axis by 3 Å. Protofilament B adopts the same structure as in AD. e, Cryo-EM density map and atomic model of triple filament (TF). Three identical protofilaments (A, B and C) extend from G273/304-E380. An additional non-proteinaceous density at the filament’s three-fold axis is labelled with a red arrow. Inset: Zoomed-in view showing one of the three identical protofilament interfaces and K331 residues from each protofilament coordinating the additional density. f, Overlay of individual protofilaments from TF with mutation V337M (orange), PHF with mutation V337M (blue) and PHF from AD (white), viewed at a 45° angle to the filaments’ axes, as in panel d.

sBy cryo-EM, we observed the presence of the Alzheimer tau fold in all three cases (Figure 1 and 2) (10). Paired helical filaments (PHFs) and straight filaments (SFs) were found in cases 1 and 3, while only PHFs were in evidence in case 2. The structures of PHFs and SFs were determined to 2.7-3.3 Å resolution and were compared to the previously determined structures of PHFs and SFs from AD (10).

PHFs from the cases with mutation V337M were nearly identical to those of PHFs assembled from wild-type tau in AD. The structures of V337M SFs and AD SFs, which comprise two asymmetrically packed protofilaments A and B with the Alzheimer tau fold, had also similar cross-sections perpendicular to the helical axis. However, unlike SFs of AD, the backbone traces of the protofilaments differed from each other along the helical axis in V337M SFs. In protofilament A, strand β4 of the Alzheimer fold, which comprises residues 336-341, was shifted along the helical axis by about 3 Å compared to protofilament B, which adopted a typical Alzheimer fold (Figure 2d). Since β4 contains the V337M mutation site, this shift may have resulted from the presence of the mutant residue. The side chain of methionine is bulkier than that of valine, but it is also more flexible.

Cryo-EM density maps and the atomic models showed that both wild-type and mutant residues could fit into the density at position 337 (Figure 2a). Analysis of the sarkosyl-insoluble fractions by mass spectrometry also showed peptides with either M337 or V337, consistent with the presence of both wild-type and mutant alleles in disease filaments (Extended Data Figure 2). By immunoblotting of sarkosyl-insoluble tau, strong bands of 60, 64 and 68 kDa, as well as a weaker band of 72 kDa, were observed, indicating the presence of all six tau isoforms in a hyperphosphorylated state (Figure 1b).

Sarkosyl-insoluble tau from case 3 with mutation V337M contained a new filament with three-fold symmetry that we named ‘triple filament’ (TF). We determined the structure of TFs to 2.3 Å resolution (Figures 1a, 2e). Unlike PHFs and SFs, which are made of two protofilaments, TFs consist of three identical protofilaments, related by C3 symmetry, with each protofilament extending from G273/304-E380. Even at 2.3 Å resolution, the side chain density at the mutation site appeared ambiguous and could accommodate either M337 or V337. A comparison of V337M TF and PHF protofilaments showed that they have similar cross-sections perpendicular to the helical axis, but that they differ by a 3 Å shift of the β4 strand of the TF along the helical axis (Figure 2f). This shift is like that between V337M SF protofilaments A and B. It is probably essential for TF formation, since the N-terminal residues of β4 contribute to the interface between protofilaments, which differ from those of PHFs and SFs. At the interface of TF protofilaments, Q336 from one protofilament intercalates between S324 and N327 of the opposite protofilament and hydrogen bonds with G326. In return, N327 of the opposite protofilament hydrogen bonds with G334 (Figure 2e). There is a large cavity along the three-fold symmetry axis of the filament, which contains a potentially negatively charged density that is coordinated by residue K331 from each protofilament. It thus appears that like SFs, whose interface contains a non-proteinaceous density between K317 and K321 from both protofilaments, TF assembly may also require external cofactors.

In addition to tau filaments with the Alzheimer fold, TMEM106B filaments were present in case 1 with the V337M mutation (Figure 1a). This individual died aged 78. The sarkosyl-insoluble fractions from the frontal cortex of cases 2 and 3, who died aged 63 and 58, were devoid of TMEM106B filaments.

### Mutation V337M increases the rate of assembly of recombinant tau(297-391)

We performed *in vitro* assembly reactions with recombinant proteins to determine if the V337M mutation in tau(297-391) influences the rate of filament assembly when compared to wild-type tau(297-391). With V337M tau(297-391), thioflavin T (ThT) fluorescence started to increase after 90 min and reached a plateau at 180 min. With wild-type tau(297-391), ThT fluorescence began to rise after 200 min and plateaued at 300 min (Figure 3a). We then proceeded to determine the cryo-EM structures of recombinant V337M tau(297-391) filaments, which revealed the presence of a majority of PHFs and a minority of quadruple helical filaments (QHFs) (Figure 3b,c). The latter, which have been described before (16), are made of two stacked PHFs held together by electrostatic interactions. The cryo-EM density at residue 337 is consistent with a methionine residue (Figure 3c). These tau filaments exhibited a cross-over length of 580 Å, whereas PHFs from human brains have cross-over lengths of 700-800 Å. Compared to V337M PHFs from human brains, the recombinant V337M PHFs differed by a slight rotation of the β-helix region with respect to the rest of the ordered core (Figure 3d).

**Figure 3.**
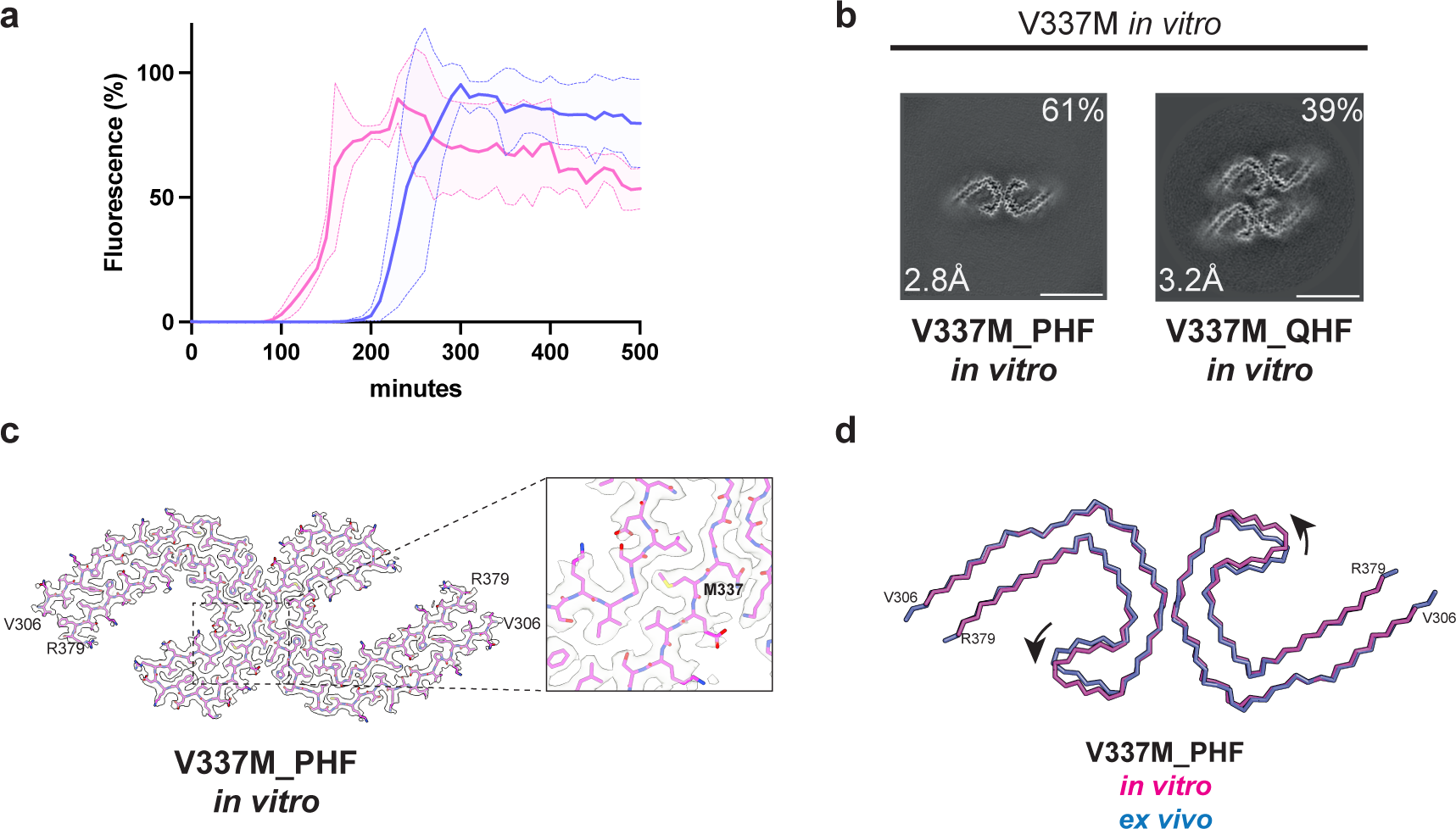
In *vitro* assembly of V337M tau(297-391). a, *In vitro* assembly assay monitored by thioflavin T (ThT) fluorescence of V337M tau(297-391) (magenta) and wild-type tau(297-391) (blue). b, Cross-sections through the cryo-EM reconstructions, perpendicular to the helical axis and with a projected thickness of approximately one rung, are shown for assembled V337M tau(297-391). c, Cryo-EM density map and atomic model of paired helical filament (PHF). Two identical protofilaments extend from V306-R379. Inset: Zoomed-in view showing the mutant methionine at position 337. d, Overlay of PHFs assembled from recombinant V337M tau(297-391) (magenta) and extracted from the frontal cortex of an individual with mutation V337M (blue). The rmsd between Cα atoms is 0.80 Å with a 9° rotation of the β-helix region relative to the rest of the ordered core being the main difference between the two structures.

### Structures of tau filaments from two cases with mutation R406W in *MAPT*

We used two previously unreported cases with mutation R406W in *MAPT*, case 1 from a US family (temporal cortex, parietal cortex and hippocampus) and case 2 from a UK family (frontal cortex, temporal cortex, parietal cortex and hippocampus). By immunohistochemistry, tau inclusions were not only present in nerve cells and their processes, but also in glial cells, chiefly astrocytes (Extended Data Figures 3-5). In the hippocampus from case 1, many extracellular tau inclusions were present, consistent with the long duration of disease. Overall, neuronal inclusions were more abundant than glial cell inclusions, which ranged from astrocytic plaques to tufted astrocytes, with many intermediates. In some brain regions from R406W case 2, reminiscent of CTE, subpial tau pathology consisting of thorn-shaped astrocytes was also present at the depths of sulci.

We determined the cryo-EM structures of tau filaments from temporal cortex, parietal cortex and hippocampus of case 1, and from frontal, temporal and parietal cortices of case 2 (Figures 4 and 5). PHFs were present in all samples, but we did not observe SFs or TFs. CTE Type I filaments were evident in temporal and parietal cortex from R406W case 2, consistent with the clinicopathological information. The structures of PHFs were determined to resolutions of 3.0-4.2 Å and found to be identical to those of AD PHFs. The ordered core of the R406W tau protofilament extended from G273/304-E380.

**Figure 4.**
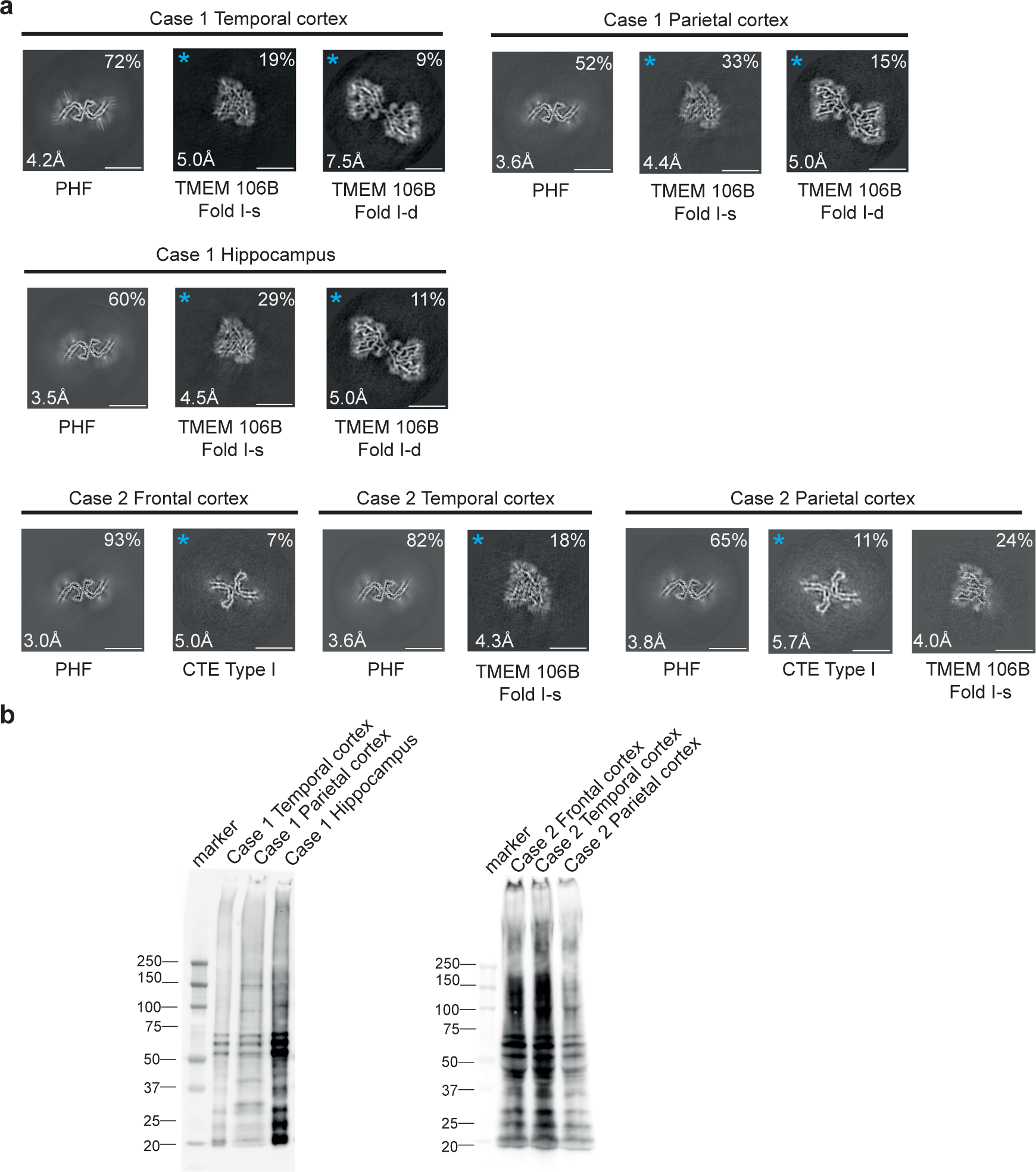
R406W mutation in *MAPT*: Cryo-EM cross-sections of tau filaments and immunoblotting. a, Cross-sections through the EM reconstructions, perpendicular to the helical axis and with a projected thickness of approximately one rung, are shown for temporal cortex, parietal cortex and hippocampus of case 1, and for frontal, temporal and parietal cortices of case 2. Resolutions (in Å) and percentages of filament types are indicated at the bottom left and top right, respectively. Scale bar, 10 nm. b, Immunoblotting of sarkosyl-insoluble tau from the temporal cortex, parietal cortex and hippocampus of case 1 with mutation R406W and from frontal, temporal and parietal cortices of case 2 with mutation R406W. Phosphorylation-independent anti-tau antibody BR134 was used.

**Figure 5.**
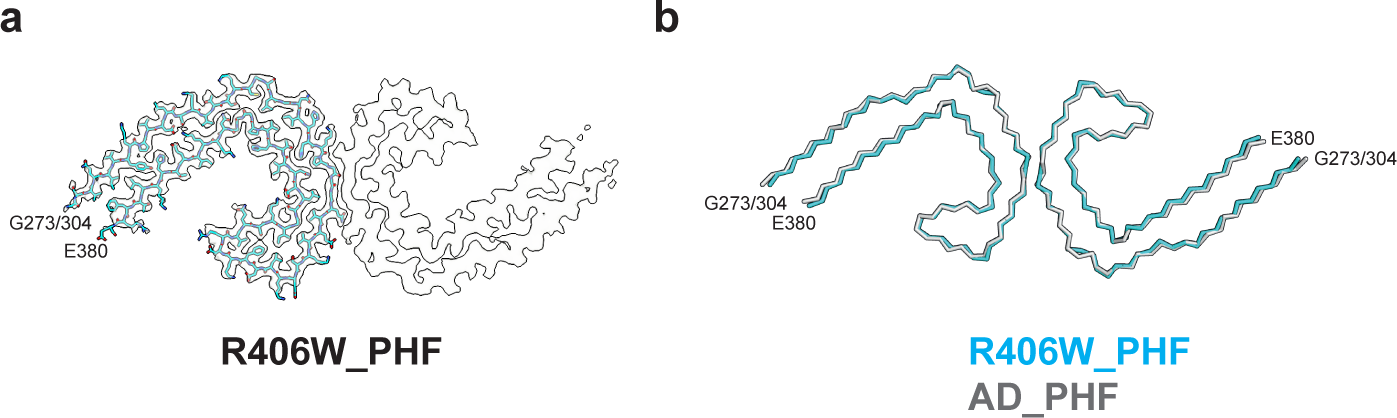
R406W mutation in *MAPT*: Cryo-EM structures of tau filaments. a, Cryo-EM density and atomic model of paired helical filament (PHF) from the frontal cortex of case 2. Two identical protofilaments extend from G273/304-E380. b, Overlay of PHFs extracted from the frontal cortex of case 2 (blue) and the frontal cortex of a case of sporadic AD (black). The rmsd between Cα atoms of the two structures is 0.78 Å.

By immunoblotting of sarkosyl-insoluble tau, we observed strong bands of 60, 64 and 68 kDa, as well as a weaker band of 72 kDa, consistent with the presence of all six tau isoforms in a hyperphosphorylated state (Figure 4b). By mass spectrometry of the sarkosyl-insoluble fractions, we detected only mutant W406 peptides, except in parietal cortex from case 2, where R406 and W406 peptides were found (Extended Data Figure 6). In addition to filaments with the Alzheimer and CTE folds, we also observed TMEM106B filaments in the sarkosyl-insoluble fractions from brain regions of both individuals, who died aged 78 and 66, respectively.

## DISCUSSION

We show that mutations V337M and R406W in *MAPT* give rise to the Alzheimer fold. V337 lies inside the ordered core of the Alzheimer fold, whereas R406 lies outside it. Small variations among the observed structures of filaments with the V337M mutation revealed the presence of an adaptable region around the mutation site that explains the accommodation of the mutant methionine without disruption of the overall fold.

By mass spectrometry, we found both wild-type and mutant tau in the core of filaments extracted from the frontal cortex of three individuals with mutation V337M, indicating that the Alzheimer tau fold can accommodate V337 and M337. Tau filaments extracted from the parietal cortex of patient 1 with mutation R406W also contained both wild-type and mutant proteins, while filaments from cerebral cortex and hippocampus of patient 2 appeared to have only mutant tau in the filaments. The former finding is in line with a study that reported the presence of wild-type and mutant forms of tau in the filaments from cases with mutation R406W (32).

Our results are consistent with positron-emission tomography (PET) scanning using [^18^F] flortaucipir that showed binding to tau inclusions in patients with *MAPT* mutations V337M and R406W (21,27,33,34). [^18^F] Flortaucipir retention has also been shown to be associated with the tau pathology of AD (35) and some prion protein amyloidoses (36). Like patients with AD, R406W mutation carriers had elevated levels of tau in cerebrospinal fluid, as measured by the antibody MTBR-tau243 (37).

The observation that cases of FTDP-17 can have the same tau filament fold as cases of AD further illustrates the fact that even though specific tau folds characterise distinct diseases, the same fold can result in clinically different conditions. Mutations in *MAPT* do not give rise to familial AD. We showed previously that other cases of FTDP-17 adopted the Pick (5) or the AGD (6) fold, depending on the relative overexpression of 3R or 4R tau.

Mutations V337M and R406W in *MAPT* led to the formation of extensive neuronal tau pathology in the form of intracellular inclusions that were reactive with antibodies RD3 and RD4, which are specific for 3R and 4R tau, respectively. In agreement with previous work (18,24), immunoblotting of sarkosyl-insoluble fractions showed a pattern of tau bands typical of all six isoforms in a hyperphosphorylated state.

In case 1 with mutation R406W, many extracellular tau inclusions (ghost tangles) were present in the hippocampus, reflecting the long duration of disease. A tangle becomes extracellular after the neuron that contained it has died. Whereas intracellular inclusions are made of full-length tau, ghost tangles progressively lose their fuzzy coat and consist mainly of the ordered filament core (R3, R4 and 10-12 amino acids after R4). These sequences are common to 3R and 4R tau isoforms. Extracellular tau inclusions can be abundant in cases with Alzheimer and CTE tau folds (38,39) and their insolubility has been attributed to extensive cross-links (40). They are much less frequent in cases with the folds of 3R and 4R tauopathies, indicating a link between filaments made of all six tau isoforms and the formation of ghost tangles.

There were also astrocytic tau inclusions in the cases with mutation R406W, suggesting that both nerve cell and glial cell inclusions contained the Alzheimer fold. Previously, shared tau folds between nerve cells and glial cells were reported for Pick’s disease, progressive supranuclear palsy, corticobasal degeneration and globular glial tauopathy (5,6,41).

It remains to be determined how mutations V337M and R406W in *MAPT* cause FTDP-17. Previous studies showed that they lead to small reductions in the ability of recombinant tau to interact with microtubules (42,43). This partial loss of function may be necessary for the assembly into filaments. It has been shown that mutations V337M and R406W do not significantly influence the heparin-induced assembly of full-length tau (44). However, the structures of heparin-induced tau filaments differ from those of AD (45). By contrast, recombinant tau(297-391) gives rise to PHFs (16). Since mutation V337M is inside the tau filament core, we assembled V337M tau(297–391); PHFs and QHFs formed, with a marked increase in the rate of filament assembly compared to wild-type tau(297-391). These findings suggest that mutation V337M has a direct effect on tau filament assembly and demonstrate the usefulness of V337M tau(297-391) for increasing filament formation in experimental studies.

## METHODS

### Cases with *MAPT* mutation V337M (Seattle family A)

We used frontal cortex from three previously described cases of Seattle family A with mutation V337M in *MAPT*. They were: Case III-1 and case IV-4 (17), as well as case IV-60 (20). We also used hippocampus from cases III-1 and IV-4. All three individuals developed a variety of symptoms, some psychiatric, consistent with a diagnosis of behavioural-variant FTD. Case 1 (III-1, UWA 63) was a female who died aged 78. At age 52 she became uncooperative, hostile, suspicious and withdrawn. She also developed progressive memory loss. Case 2 (IV-4, UWA 271) was a female who died aged 63, following an 11-year history of FTD. She was the daughter of case 1 and presented with antisocial and impulsive behaviours, which were followed by apathy, loss of language and dementia. Case 3 (IV-60, UWA 578) was a male who died at age 58 following a 16-year history of FTD. He lost his job because of poor performance and was vague, restless and behaved inappropriately. Early on, he had mild memory problems and deficient executive function. His condition slowly progressed to dementia requiring hospitalisation.

### Case with *MAPT* mutation R406W (US family)

We used temporal and parietal cortex, as well as hippocampus from a female with mutation R406W in *MAPT* who died aged 78, after a 29-year history of personality changes and cognitive impairment. The clinical diagnosis was AD. Genetic or neuropathological information on the parents was not available, but the mother had been diagnosed with AD. Besides the proband, she had four other children (three females and one male), who all developed cognitive impairment in mid-life. They were diagnosed with AD (three females) or FTDP-17 (male). The symptoms of the proband were dominated by progressive dementia and personality changes characterised by an anxiety disorder.

### Case with *MAPT* mutation R406W (UK family)

We used frontal, temporal and parietal cortices from a male with mutation R406W in *MAPT* who died aged 66 after a 9-year history of FTD. Both parents died without known FTD before the age of 61. At least one sibling developed FTD. The initial symptoms were episodic memory impairment with subsequent executive dysfunction and personality changes characterised by impulsivity and inappropriate behaviour. Magnetic resonance imaging (MRI) showed severe bilateral frontal lobe and medial temporal lobe atrophy that was more severe on the left side. This individual worked as an electrician until the age of 55 and had a history of alcohol abuse. In his youth, he had played soccer for several years.

### DNA sequencing

Genomic DNA was extracted from blood with informed consent. Standard amplification reactions were done with 50 ng genomic DNA, followed by DNA sequencing of exons 1 and 9-13 of *MAPT* with adjoining intronic sequences, as described (46).

### Filament extraction from human brains

Sarkosyl-insoluble material was extracted from frontal cortex of cases 1-3 with mutation V337M and frontal, temporal and parietal cortices of two cases with mutation R406W in *MAPT*, as described (47). Hippocampus from case 2 was also used. Tissues were homogenised in 20 vol (w/v) buffer A (10 mM Tris-HCl, pH 7.4, 0.8 M NaCl, 10% sucrose and 1 mM EGTA), brought to 2% sarkosyl and incubated at 37° C for 30 min. The samples were centrifuged at 7,000 g for 10 min, followed by spinning of the supernatants at 100,000 g for 60 min. The pellets were resuspended in buffer A (1 ml/g tissue) and centrifuged at 9,500 g for 10 min. The supernatants were diluted 3-fold in buffer B (50 mM Tris-HCl, pH 7.5, 0.15 M NaCl, 10% sucrose and 0.2% sarkosyl), followed by a 60 min spin at 100,000 g. For cryo-EM, the pellets were resuspended in 100 μl/g buffer C (20 mM Tris-HCl, pH 7.4, 100 mM NaCl).

### Immunoblotting and histology

For immunoblotting, samples were resolved on 4-12% Bis-Tris gels (NuPage) and the primary antibody [BR134 (48), 1:1,000] was diluted in PBS plus 0.1% Tween 20 and 5% non-fat dry milk. Histology and immunohistochemistry were carried out as described (46). Some sections (8 μm) were counterstained with haematoxylin. The primary antibodies were AT8 (Thermo-Fisher; 1:1,000 or 1:300), RD3 (Sigma-Millipore, 1:3,000) RD4 (Sigma-Millipore 1:100) and anti-4R (Cosmo Bio 1:400).

### Expression and purification of recombinant tau(297-391) with and without mutation V337M

Tau(297-391) with the V337M mutation was made using *in vivo* assembly (49). Reverse and forward primers were designed to share 15 nucleotides of homologous region and 15-30 nucleotides for annealing to the template. Expression of tau(297-391) was carried out in *E. coli* BL21(DE3)-gold cells (Agilent Technologies), as described (50). One plate of cells was resuspended in 1 L 2xTY (tryptone yeast) supplemented with 2.5 mM MgSO_4_ and 100 mg/L ampicillin and cells were grown to an optical density of 0.8 at 600 nm. They were induced by the addition of 1 mM IPTG for 4 h at 37° C, collected by centrifugation (4,000g for 30 min at 4° C) and flash frozen. The pellets were resuspended in washing buffer at room temperature (50 mM MES, pH 6.5, 10 mM EDTA, 10 mM DTT, 0.1 mM PMSF) and cells were lysed by sonication (90% amplitude using a Sonics VCX-750 Vibracell Ultra Sonic Processor for 4 min, 3s on/6s off) at 4° C. The lysed cells were centrifuged at 20,000g for 35 min at 4° C, filtered through 0.45 μm cut-off filters and loaded onto a HiTrap CaptoS 5-ml column at 4° C (GE Healthcare). The column was washed with 10 vol. of buffer, followed by elution through a gradient of washing buffer containing 0-1M NaCl. Fractions of 3.5 ml were collected and analysed by SDS-PAGE (4-20% Tris-glycine gels). Protein-containing fractions were pooled and precipitated using 0.3 g/ml ammonium sulphate and left on a rocker for 30 min at 4° C. The solution was then centrifuged at 20,000g for 35 min at 4° C and resuspended in 2 ml of 10 mM potassium phosphate buffer, pH 7.2, containing 10 mM DTT, and loaded onto a 16/600 75-pg size-exclusion column. Fractions were analysed by SDS-PAGE and protein-containing fractions pooled and concentrated at 4° C to 20 mg/ml using molecular weight concentrators with a cut-off filter of 3kDa. Purified protein samples were flash-frozen in 50-100 μl aliquots. Protein concentrations were determined using a NanoDrop2000 (Thermo Fisher Scientific).

### Filament assembly of recombinant tau(297-391) with and without mutation V337M

Prior to assembly, proteins and buffers were filtered through sterile 0.22 μM Eppendorf filters. A solution of 6 mg/ml wild-type tau(297-391) or V337M tau(297-391) was prepared at room temperature in 50 mM KPOH, pH 7.2, 10 mM DTT and 2 μM Thioflavin T (ThT). An additional set of reactions was prepared without ThT for cryo-EM analysis. Thirty μl aliquots were dispensed in a 384-well plate (company) that was sealed and placed in a Fluostar Omega (BMG Labtech) plate reader. Assembly was carried out using orbital shaking (200 rpm) at 37° C for 12 h.

### Mass spectrometry

Sarkosyl-insoluble pellets were resuspended in 200 ml hexafluoroisopropanol. Following a 3 min sonication at 50% amplitude (QSonica), they were incubated at 37° C for 2 h and centrifuged at 100,000 g for 15 min, before being dried by vacuum centrifugation. Protein samples resuspended in 4M urea, 50 mM ammonium bicarbonate (ambic) were reduced with 5mM DTT at 37°C for 40 min and alkylated with 10 mM chloroacetamide for 30 min. For V337M samples, they were digested with LysC (Promega) for 4 h, followed by trypsin after dilution of urea to 1.5 M. For R406W samples, urea was diluted to 1.0 M and incubated with AspN (Promega) overnight at 30°C. Digestion was stopped by the addition of formic acid to a final concentration of 0.5%, followed by a centrifugation at 16,000 g for 5 min. The supernatants were desalted and fractionated using home-made C18 stage tips (3M Empore) packed with poros oligo R3 (Thermo Scientific) resin. Bound peptides were eluted stepwise with increasing MeCN in 10 mM ambic and partially dried in a SpeedVac (Savant). Samples were analysed by LC-MS/MS using a Q Exactive Plus hybrid quadrupole-Orbitrap mass spectrometer (Thermo Fisher Scientific) coupled online to a fully automated Ultimate 3,000 RSLC nano System (Thermo Scientific). LC-MS/MS data were searched against the human reviewed database (UniProt, downloaded 2023), using the Mascot search engine (Matrix Science, v.2.80. Scaffold (version 4, Proteome Software Inc.) was used to validate MS/MS-based peptide and protein identifications.

### Electron cryo-microscopy

Cryo-EM grids (Quantifoil 1.2/1.3, 300 mesh) were glow-discharged for 1 min using an Edwards (S150B) sputter coater. Three μl of the sarkosyl-insoluble fractions or recombinant Tau assemblies were applied to the glow-discharged grids, followed by blotting with filter paper and plunge freezing into liquid ethane using a Vitrobot Mark IV (Thermo Fisher Scientific) at 4° C and 100% humidity. Cryo-EM images were acquired on a Titan Krios G2 or G4 microscope (Thermo Fisher Scientific) operated at 300 kV and equipped with a Falcon-4 or a Falcon-4i direct electron detector. Images were recorded for 2s in electron event representation format (51), with a total dose of 40 electrons per A^2^ and a pixel size of 0.824 Å (Falcon-4) or 0.727 Å (Falcon-4i). See Extended Data Table 1 and Extended Data Figure 7,8 for further details.

### Data processing

Datasets were processed in RELION using standard helical reconstruction (52,53). Movie frames were gain-corrected, aligned and dose-weighted using RELION’s own motion correction programme (54). Contrast transfer function (CTF) was estimated using CTFFIND4.1 (55). Filaments were picked manually and segments were extracted with a box size of 1,024 pixels, prior to downsizing to 256 pixels. Reference-free 2D classification was carried out and selected class averages were re-extracted using a box size of 400 pixels. Initial models were generated *de novo* from 2D class average images using relion_helix_inimodel2d (56). Three-dimensional refinements were performed in RELION-4.0 and the helical twist and rise refined using local searches. Bayesian polishing and CTF refinement were used to further improve resolutions (57). The final maps were sharpened using post-processing procedures in RELION-4.0 and resolution estimates were calculated based on the Fourier shell correlation (FSC) between two independently refined half-maps at 0.143 (Extended Data Figure 8) (58). We used relion_helix_toolbox to impose helical symmetry on the post-processing maps.

### Model building and refinement

Atomic models were built manually using Coot (59), based on published structures (PHF, PDB:5O3L; SF, PDB:5O3T) (10). Model refinements were performed using ISOLDE (60), *Servalcat* (61) and REFMAC5 (62,63). Models were validated with MolProbity (64). Figures were prepared with ChimeraX (65) and PyMOL (66).

## Acknowledgements

We thank the patients’ families for donating brain tissues, T. Darling, I. Clayson and J. Grimmett for help with high-performance computing and the EM facility of the Medical Research Council (MRC) Laboratory of Molecular Biology for help with cryo-EM data acquisition. We are grateful to R. Richardson, N. Maynard, M. Jacobsen and B. Glazier for help with histology and immunohistochemistry. This work was supported by the MRC, as part of U.K. Research and Innovation (MC_UP_A025_1013 to S.H.W.S. and MC_1051284291 to M.G.). It was also supported by the U.S. National Institutes of Health (P30-AG010133, R01-AG080001 and RF1-AG071177, to B.G.) and the Department of Pathology and Laboratory Medicine, Indiana University School of Medicine (to B.G.). The Queen Square Brain Bank is supported by the Rita Lila Weston Institute for Neurological Studies.

## Author contributions

M.B., C.L., J.R.M., P.W.C., Z.J., T.D.B. and B.G. identified patients and performed neuropathology and DNA sequencing. C.Q. performed immunoblot analysis. C.Q., S.P.-C and C.F. performed mass spectrometry. C.Q. and S.L. collected cryo-EM data. C.Q., S.L., A.G.M. and S.H.W.S. analysed cryo-EM data. S.H.W.S. and M.G. supervised the project. All authors contributed to the writing of the manuscript.

## Data availability

Cryo-EM maps have been deposited in the Electron Microscopy Data Bank (EMDB) with accession numbers: EMD-19846; EMD-19849; EMD-19852; EMD-19854; EMD-19855. Corresponding refined atomic models have been deposited in the Protein Data Bank (PDB) under the following accession numbers: 9EO7; 9EO9; 9EOE; 9EOG; 9EOH. Please address requests for materials to the corresponding authors.

---

**Extended Data Figure 1.**
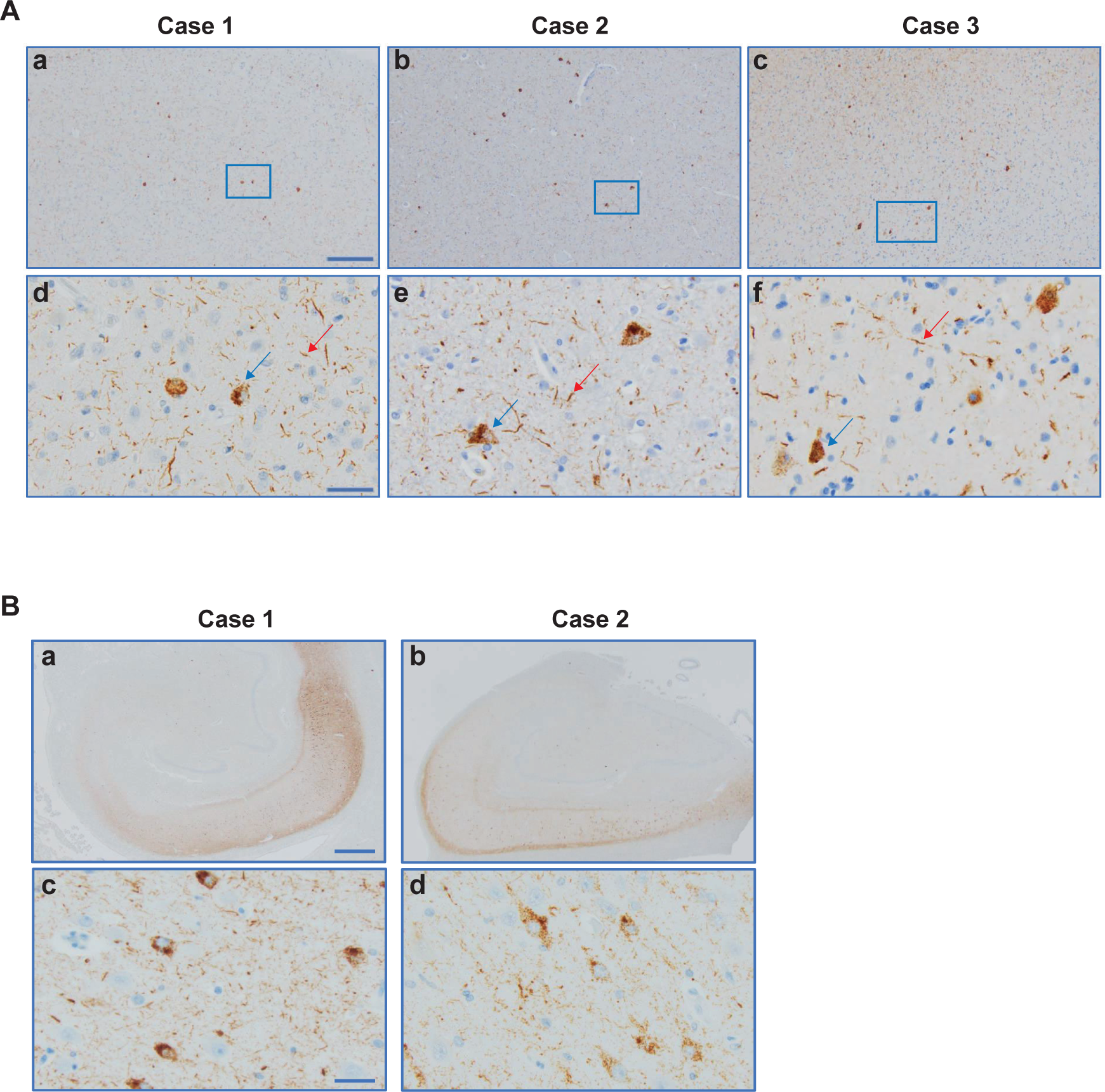
V337M mutation in *MAPT*: Immunohistochemical localisation of tau inclusions in frontal cortex and hippocampus. (a-f) Tau pathology in grey matter. Higher magnifications of tissue areas within the insets in a-c are shown in d-f. Intraneuronal inclusions (blue arrows) and neuropil threads (red arrows) are in evidence. Antibody: AT8. Scale bars: 200 μm (a-c); 40 μm (d-f). a,b, Low-power view; c,d, neurofibrillary tangles and neuropil threads in the pyramidal cell layer. Antibody: AT8. Scale bars: 1,000 μm (a,b) and 40 μm (c,d).

**Extended Data Figure 2.**
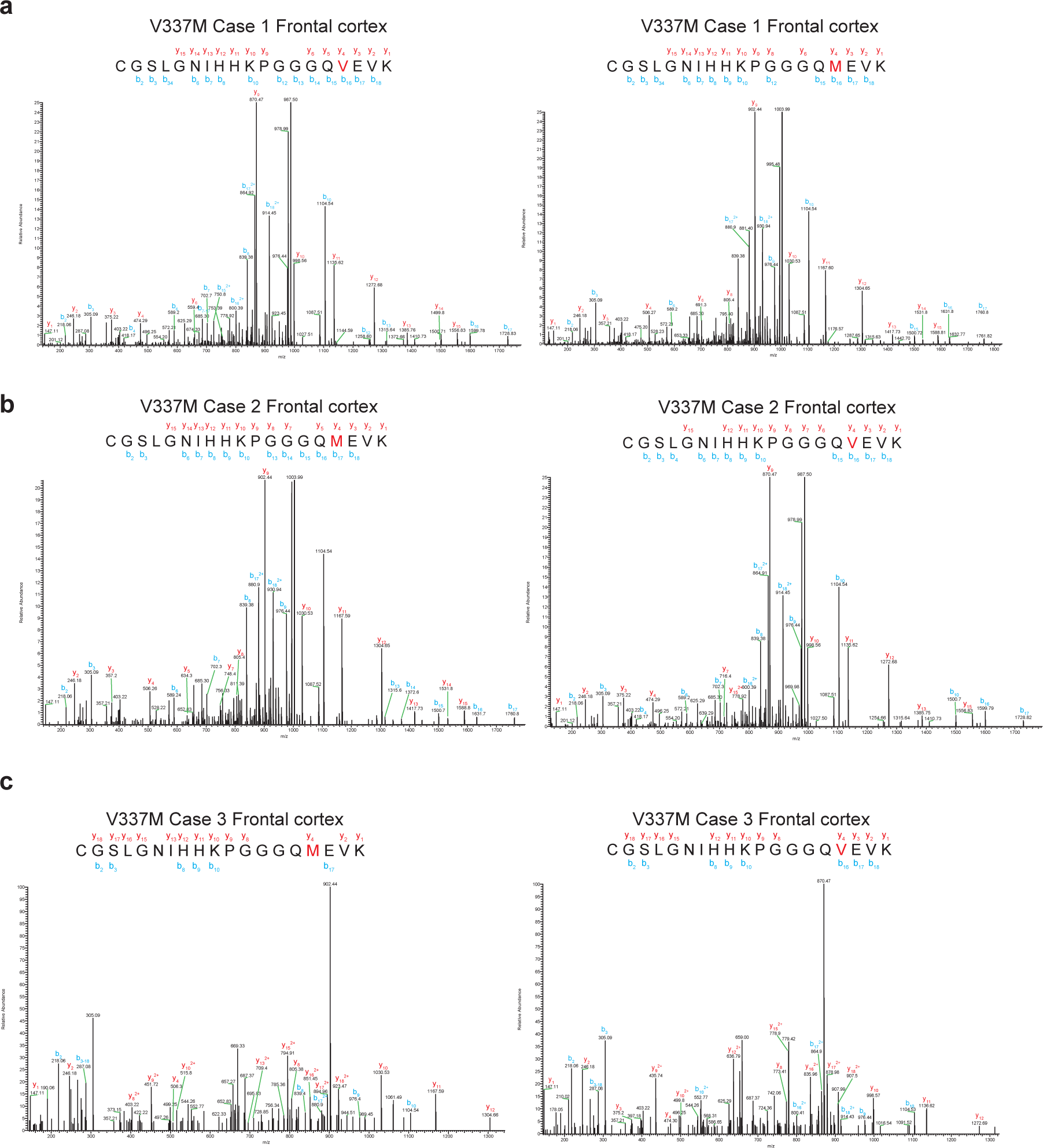
Mass spectrometry of tau from the sarkosyl-insoluble fractions of cases with mutation V337M in *MAPT*. MALDI mass spectra of the frontal cortex from cases 1-3 (a-c). Wild-type (V337) and mutant (M337) peptides were detected.

**Extended Data Figure 3.**
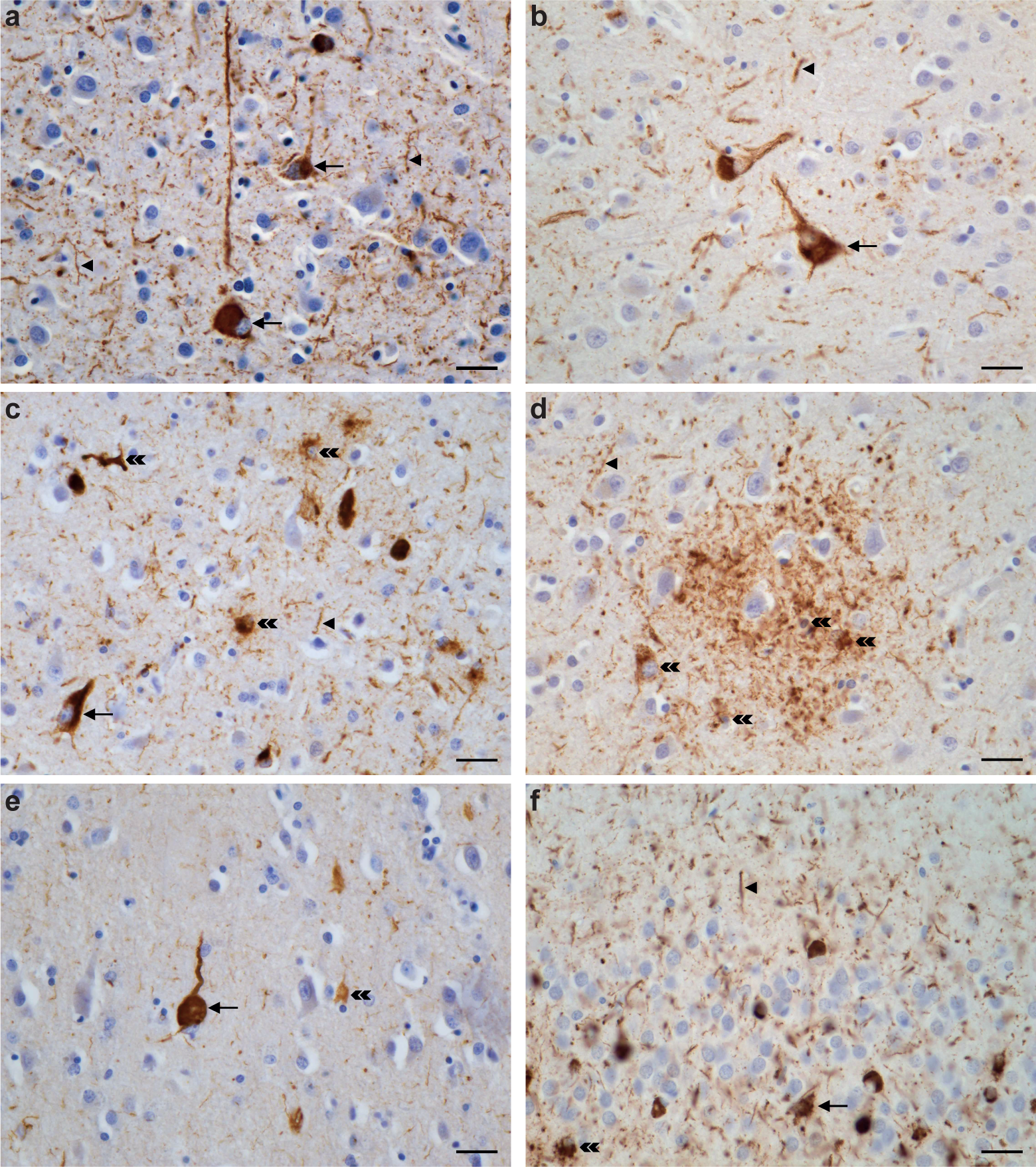
R406W mutation in *MAPT*: Immunohistochemical localisation of tau in temporal cortex, parietal cortex and hippocampus of case 1. a,c,e, Tau-immunopositive nerve cell bodies (arrows) and neuropil threads (arrowheads) are shown in temporal cortex; b,d, parietal cortex; f, hippocampus. c,d,e,f, Labelled astrocytes (double arrowheads). Panel (d) shows plaques composed of numerous threads, corresponding probably to the processes of neurons and astrocytes. Antibodies: AT8 (a,b,d,f); RD4 (c); RD3 (e). Scale bar, 25 μm.

**Extended Data Figure 4.**
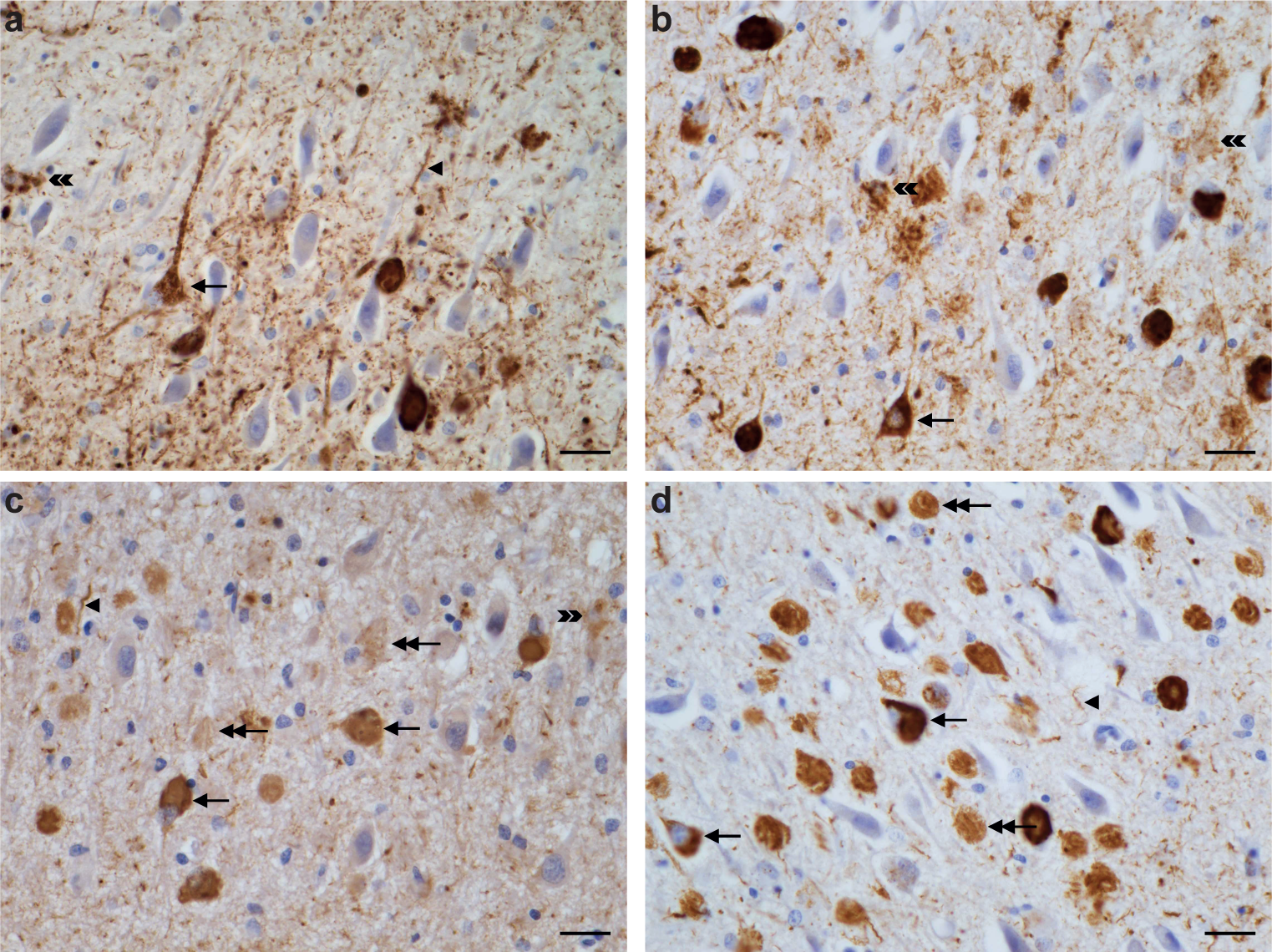
R406W mutation in *MAPT*: Immunohistochemical localisation of tau inclusions in the hippocampus of case 1. The pyramidal layer is shown. Tau-immunopositive intracellular neuronal inclusions (single headed arrows) and extracellular ghost inclusions (double headed arrows), as well as neuropil threads (single arrowheads) and astrocytic inclusions (double arrowheads) are indicated. Antibodies: AT8 (a), RD4 (b), anti-4R (c), RD3 (d). Scale bar, 25 μm.

**Extended Data Figure 5.**
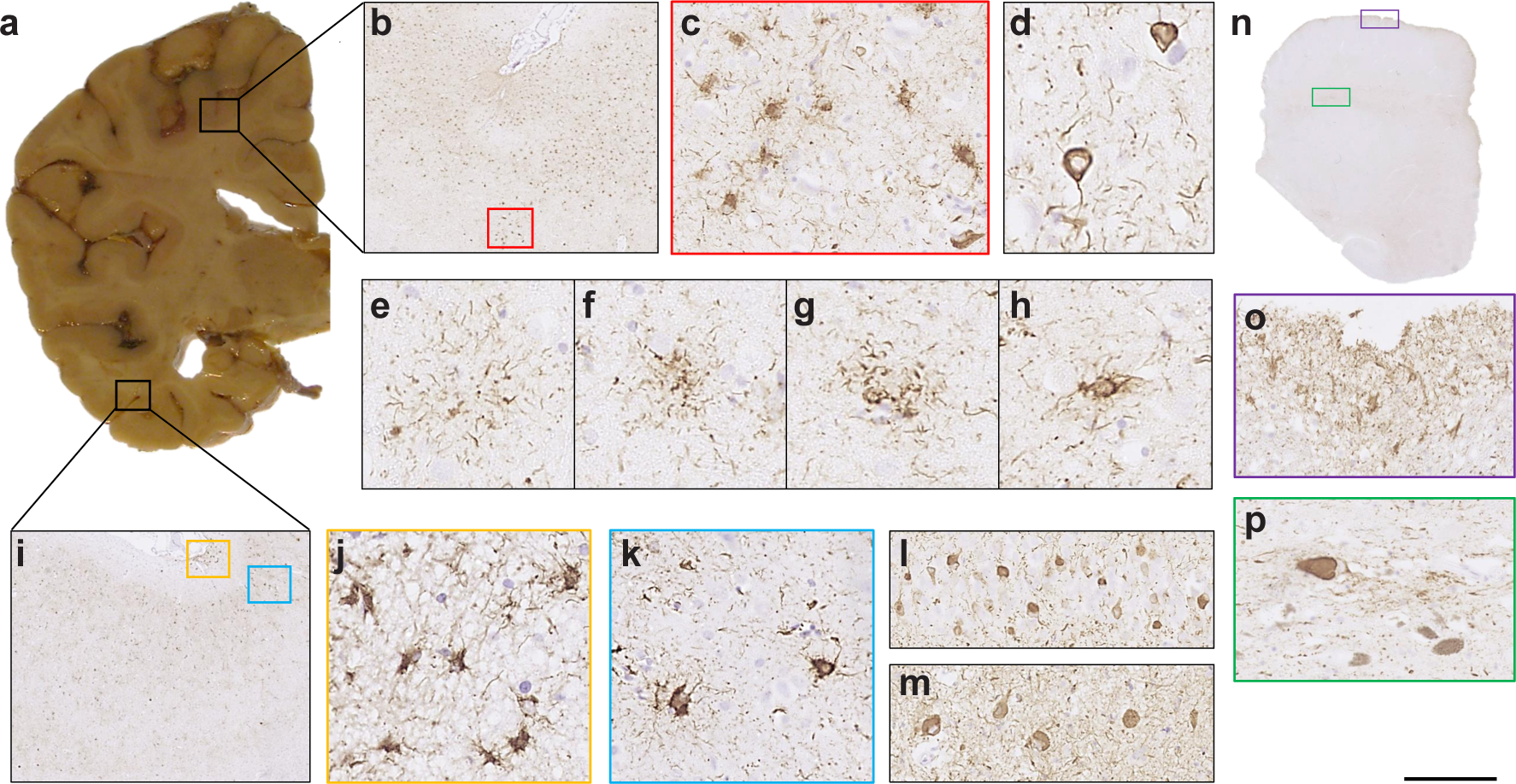
R406W mutation in *MAPT*: Immunohistochemical localisation of tau inclusions in case 2. a, Mild atrophy of the frontal cortex, severe atrophy of the temporal cortex and underlying white matter, with marked reduction in bulk of the hippocampus. Anterior and temporal horns of the lateral ventricle are dilated; b, tau pathology in the anterior frontal cortex; c, some stained cells resemble tufted or thorn-shaped astrocytes; d, abundant neuronal inclusions and neuropil threads; e, astrocytic plaque; f,g, structures in-between astrocytic plaques and tufted astrocytes; h, tufted astrocyte; i,k, Tau pathology in the lateral temporal cortex was similar to that in the anterior frontal cortex; l, CA4 region of the hippocampus; m, dentate gyrus; j, subpial astrocytic tau pathology at the depth of a sulcus in the lateral temporal cortex. n, Low-power view of the midbrain; o, subpial astrocytic tau pathology; p, neuronal tau staining in the substantia nigra. AT8 antibody. Scale bar: b, 750 μm; c,k, 70 μm; d, 40 μm; e,f,g,h, 30 μm; i, 670 μm; j, 50 μm; l. 100 μm; m, 110 μm; n, 5.5 mm; o,p, 90 μm.

**Extended Data Figure 6.**
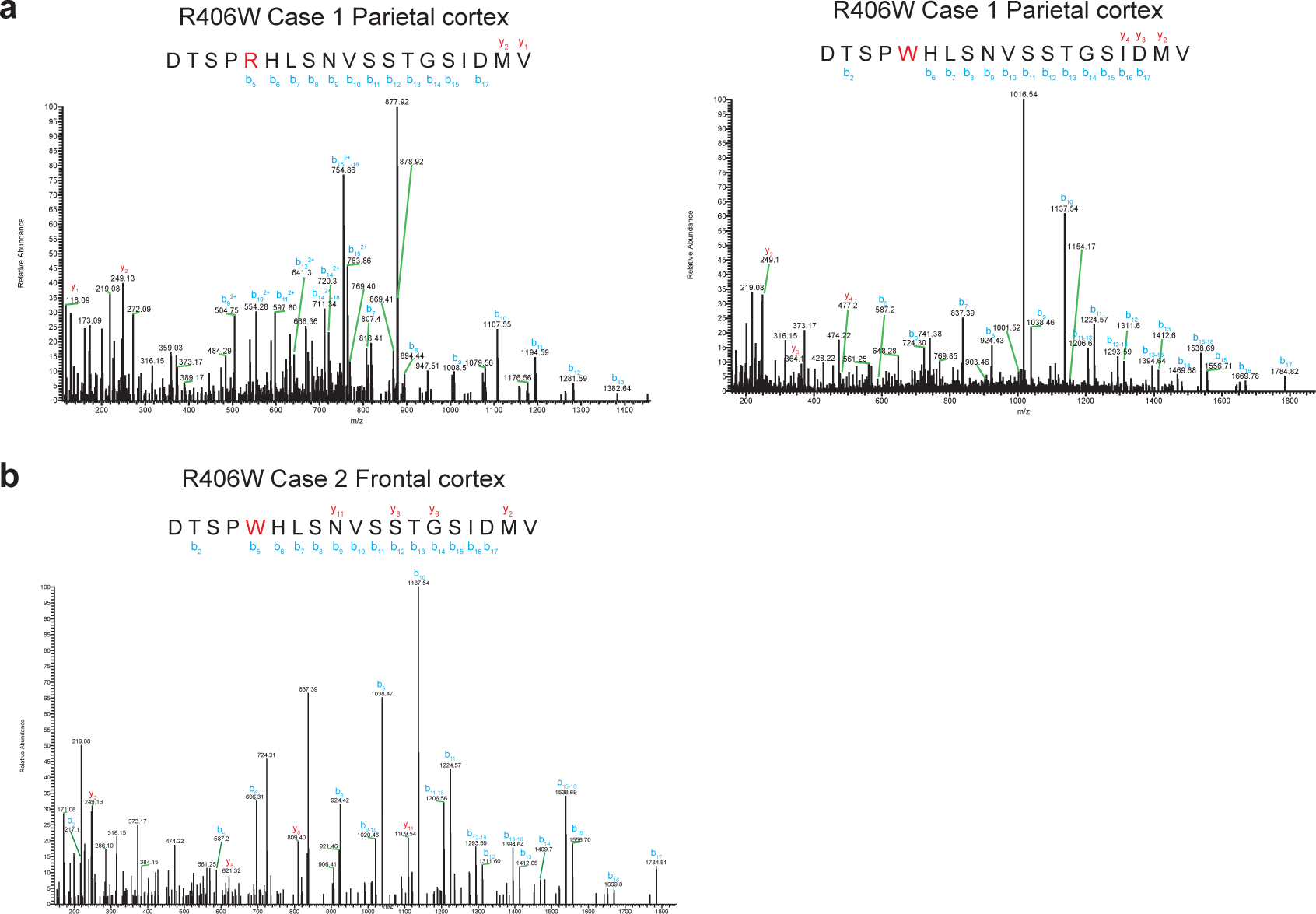
Representative mass spectrometry of tau from the sarkosyl-insoluble fractions of cases with mutation R406W in *MAPT*. MALDI mass spectra of parietal cortex from case 1 (a) and frontal cortex from case 2 (b). Wild-type (R406) and mutant (W406) peptides were detected in parietal cortex from case 1. Only mutant peptides (W406) were detected in frontal cortex from case 2. Similarly, only mutant peptides were detected in temporal cortex and hippocampus from case 1, as well as in temporal cortex, parietal cortex and hippocampus from case 2.

**Extended Data Figure 7.**
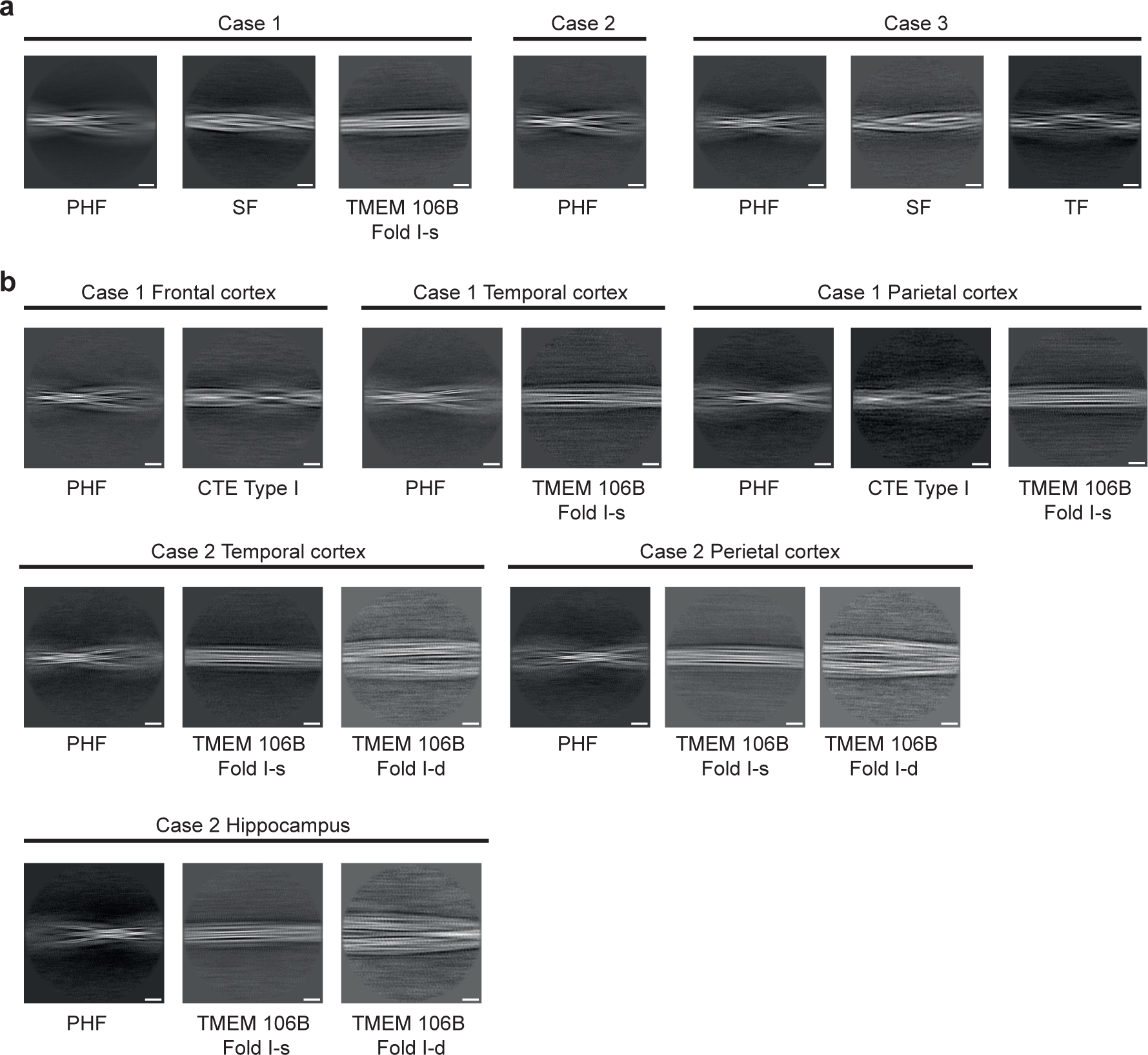
Two-dimensional class averages of filaments from three cases with mutation V337M (a) and two cases with mutation R406W (b) in *MAPT*. Scale bar, 10 nm.

**Extended Data Figure 8.**
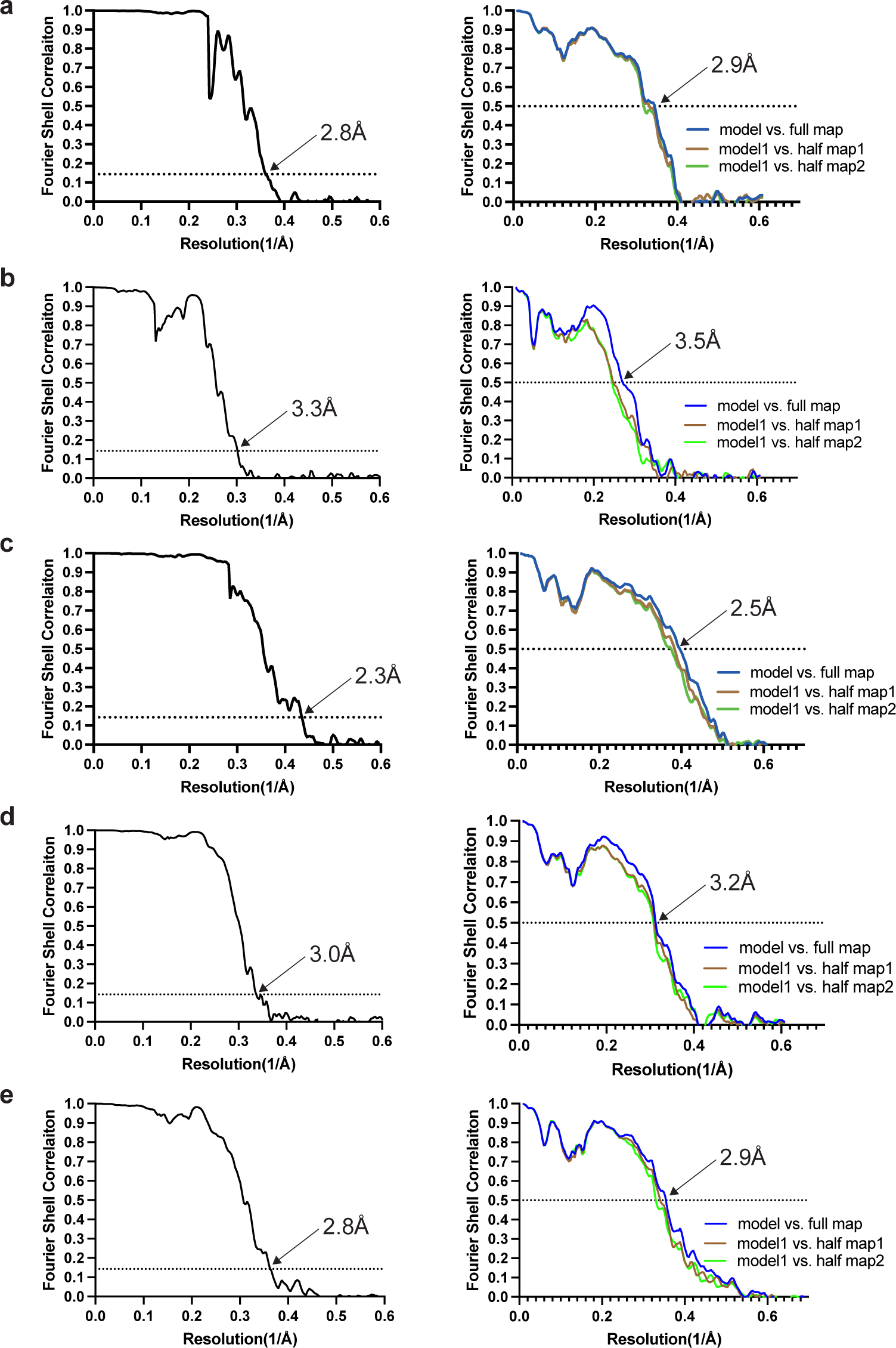
Fourier shell correlation (FSC) curves. FSC curves of cryo-EM maps (left panel) and model to map validation (right panel). a, paired helical tau filament from V337M mutant; b, straight tau filament from V337M mutant; c, triple tau filament from V337M mutant; d, paired helical tau filament from R406W mutant; e, paired helical tau filament from assembled V337M tau (297-391).

**Extended Data Table 1:**
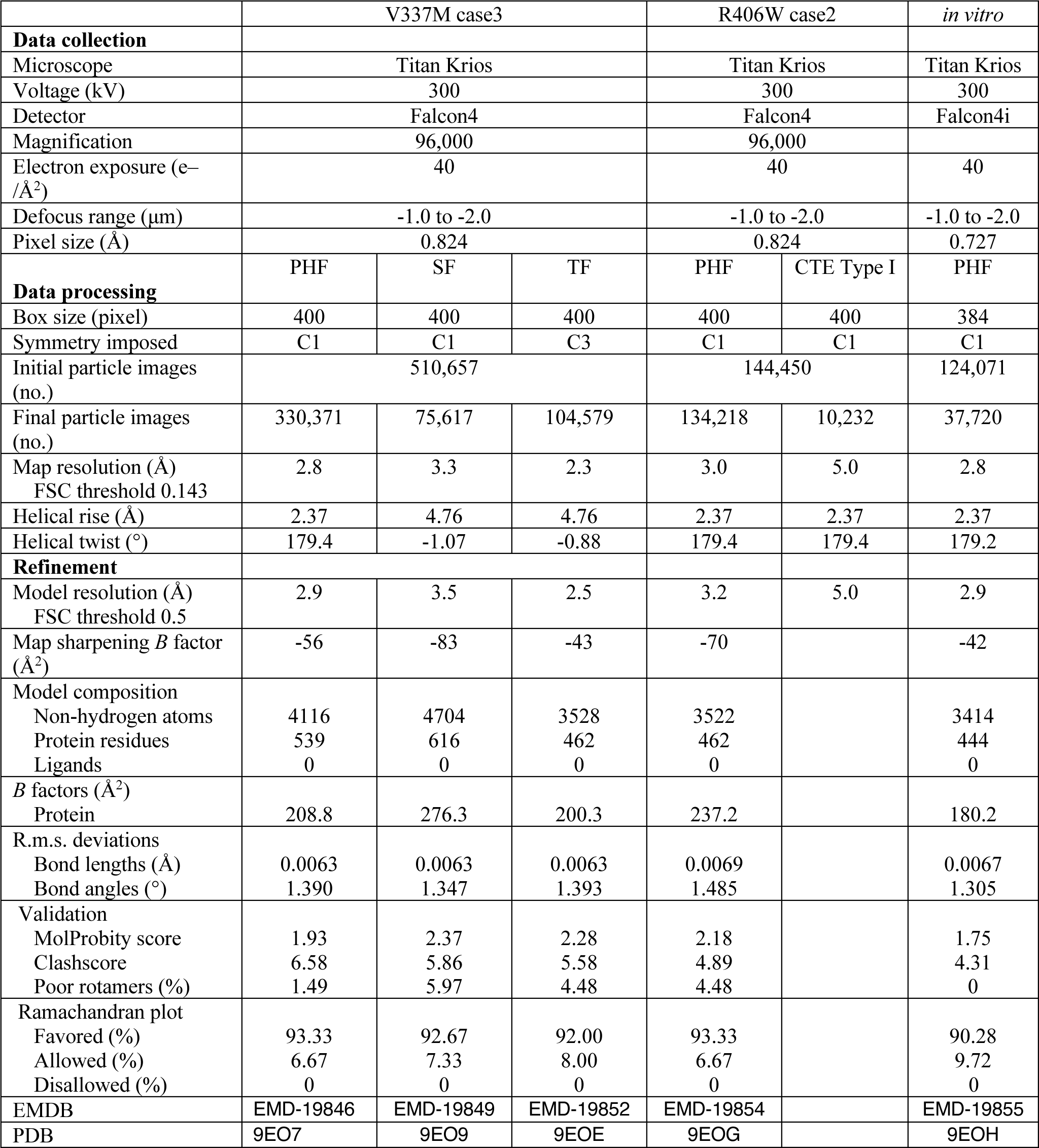
Cryo-EM data collection, refinement and validation statistics.

|  | V337M case3 |  |  | R406W case2 |  | *in vitro* |
| --- | --- | --- | --- | --- | --- | --- |
| **Data collection** |  |  |  |  |  |  |
| Microscope | Titan Krios |  |  | Titan Krios |  | Titan Krios |
| Voltage (kV) | 300 |  |  | 300 |  | 300 |
| Detector | Falcon4 |  |  | Falcon4 |  | Falcon4i |
| Magnification | 96,000 |  |  | 96,000 |  |  |
| Electron exposure (e<sup>-</sup>/Å<sup>2</sup>) | 40 |  |  | 40 |  | 40 |
| Defocus range (μm) | -1.0 to -2.0 |  |  | -1.0 to -2.0 |  | -1.0 to -2.0 |
| Pixel size (Å) | 0.824 |  |  | 0.824 |  | 0.727 |
| **Data processing** | PHF | SF | TF | PHF | CTE Type I | PHF |
| Box size (pixel) | 400 | 400 | 400 | 400 | 400 | 384 |
| Symmetry imposed | C1 | C1 | C3 | C1 | C1 | C1 |
| Initial particle images (no.) | 510,657 |  |  | 144,450 |  | 124,071 |
| Final particle images (no.) | 330,371 | 75,617 | 104,579 | 134,218 | 10,232 | 37,720 |
| Map resolution (Å)<br>FSC threshold 0.143 | 2.8 | 3.3 | 2.3 | 3.0 | 5.0 | 2.8 |
| Helical rise (Å) | 2.37 | 4.76 | 4.76 | 2.37 | 2.37 | 2.37 |
| Helical twist (°) | 179.4 | -1.07 | -0.88 | 179.4 | 179.4 | 179.2 |
| **Refinement** |  |  |  |  |  |  |
| Model resolution (Å)<br>FSC threshold 0.5 | 2.9 | 3.5 | 2.5 | 3.2 | 5.0 | 2.9 |
| Map sharpening *B* factor (Å<sup>2</sup>) | -56 | -83 | -43 | -70 |  | -42 |
| Model composition |  |  |  |  |  |  |
| Non-hydrogen atoms | 4116 | 4704 | 3528 | 3522 |  | 3414 |
| Protein residues | 539 | 616 | 462 | 462 |  | 444 |
| Ligands | 0 | 0 | 0 | 0 |  | 0 |
| *B* factors (Å<sup>2</sup>) |  |  |  |  |  |  |
| Protein | 208.8 | 276.3 | 200.3 | 237.2 |  | 180.2 |
| R.m.s. deviations |  |  |  |  |  |  |
| Bond lengths (Å) | 0.0063 | 0.0063 | 0.0063 | 0.0069 |  | 0.0067 |
| Bond angles (°) | 1.390 | 1.347 | 1.393 | 1.485 |  | 1.305 |
| Validation |  |  |  |  |  |  |
| MolProbity score | 1.93 | 2.37 | 2.28 | 2.18 |  | 1.75 |
| Clashscore | 6.58 | 5.86 | 5.58 | 4.89 |  | 4.31 |
| Poor rotamers (%) | 1.49 | 5.97 | 4.48 | 4.48 |  | 0 |
| Ramachandran plot |  |  |  |  |  |  |
| Favored (%) | 93.33 | 92.67 | 92.00 | 93.33 |  | 90.28 |
| Allowed (%) | 6.67 | 7.33 | 8.00 | 6.67 |  | 9.72 |
| Disallowed (%) | 0 | 0 | 0 | 0 |  | 0 |
| EMDB | EMD-19846 | EMD-19849 | EMD-19852 | EMD-19854 |  | EMD-19855 |
| PDB | 9EO7 | 9EO9 | 9EOE | 9EOG |  | 9EOH |

